# The Genetic Architecture of Neonatal Deer Mouse Cries Implicates the Cerebellum in the Temporal Control of an Infant Social Behavior

**DOI:** 10.64898/2026.09.18.752619

**Authors:** N Jourjine, ML Woolfolk, HE Hoekstra

## Abstract

Vocal communication is a conserved vertebrate social behavior that begins at birth with the cries of infants. These neonatal vocal signals vary across contexts to elicit parental care that matches infant needs, and across species to match species-specific ecologies and social systems. This natural variation presents an opportunity to identify proximate mechanisms supporting flexible neonatal vocal behaviors and their evolution. To this end, we recently described neonatal vocalizations in North American deer mice, a model system for behavioral evolution, and identified heritable interspecific differences in features of these vocalizations. Here, we examine one of these features, temporal structure, which in humans and rodents contains information about infant distress. We find that temporal features of neonatal deer mouse cries have diverged more across deer mouse species than spectral features, and that in a playback assay cry duration affects the ability of pup vocalizations to elicit parental approach. To test proximate mechanisms underlying natural variation in cry duration, we first focus on the cerebellar system, a conserved hindbrain region that contributes to the temporal structure of motor rhythms. We identify interspecies variation in the gross anatomy of the neonatal cerebellum and inferior olive and show that pharmacological perturbation of olivocerebellar function alters the temporal structure of neonatal cries in deer mice. Next, we use an interspecies cross to map the genetic architecture underlying neonatal cry duration in deer mice. We identify a single quantitative trait locus significantly associated with cry duration, as well as candidate genes in this locus that could plausibly underlie species differences in the temporal structure of infant crying. Among these candidates are genes that are differentially expressed between species, function in the developing cerebellum, and have been linked to the duration of neonatal vocal signals in house mice. Taken as a whole, this work identifies contributions of the cerebellar system to neonatal social behaviors and suggests testable neural and genetic hypotheses about the evolution of infant crying.

## Introduction

In mammals, behavioral interactions between parents and infants are critical for early-life survival. Distantly related species share core elements of these behaviors, suggesting conserved underlying mechanisms^1^. At the same time, specific features of parent-offspring social behaviors can be highly variable even between closely related species^2^. This pattern of diversification raises fundamental questions about the neural basis of sociality and its evolution: What neurophysiological mechanisms support adaptive parent-offspring social interactions within species? How have these mechanisms evolved to produce the diversity of parent-offspring social dynamics observed across species? A large body of work has addressed these questions by identifying proximate mechanisms underlying infant-directed parental care behaviors and their evolution^3–6^. The parent-infant social dynamic also critically involves the social behaviors of infants themselves^7^, yet the proximate underpinnings of parent-directed infant behaviors have received relatively less attention. As a result, the mechanisms underlying infant social behaviors and their evolution represent a gap in our understanding of parent-offspring sociality.

Parent-directed vocalization is one deeply conserved infant social behavior that is critical for early survival across vertebrates. In mammals, infant vocalizations often begin at the moment of birth, where they trigger diverse forms of parental care required for neonatal health. Vocal signals elicit this care via a set of features that appear to be at least partially conserved across species^8^. These include the presence of low-frequency “cries” with atonal or chaotic elements^9^, the presence of rhythmic vocal bouts^10^, and the importance of temporal features, like call duration, in eliciting parental attention^11,12^. Infants also exhibit extensive variation in the temporal and spectral parameters they apply to this conserved framework. Within species, this variation depends on physiological need, social context, and stage of postnatal development, among other factors. Infant crying behaviors also vary between species, with some evidence that crying has been a target of natural selection imposed by species-specific social systems. For example, the cries of infant Cape fur seals (*Arctocephalus pusillus pusillus*) exhibit an unusually high degree of vocal stereotypy^13^ compared to closely related species, possibly^14^ an adaptation to a unique rearing environment in which mothers must regularly identify their own pups’ calls in groups of hundreds of vocalizing individuals. However, the proximate mechanisms underlying species-specific infant vocal features such as this remain unclear.

One likely contributor to these features is the neural architecture underlying vocal production. In adults, vocalization is controlled by a core neural circuit consisting of the midbrain periaqueductal gray (PAG), which integrates cortical inputs to gate vocalization through hindbrain nuclei such as the nucleus ambiguus (RAm). These in turn drive laryngeal motor neurons in coordination with breathing^15^. While this pathway was first elucidated using focal lesioning and electrical stimulation in primates, more recent studies have used cell-type specific perturbations to demonstrate that its central features are conserved in adult rodents^16–20^, arguing that it reflects a neural architecture widely shared across mammals. Rodent studies have further extended our understanding of mammalian vocal production by, for example, linking species divergence in cortical^21^ and PAG^22^ circuits to the evolution of vocal repertoires, and identifying potentially novel components of the core vocal production pathway, such as a recently described cerebellar-PAG circuit that is sufficient to gate adult vocalization^23^.

Rodent studies have also extended our understanding of neonatal vocalization, where most work has focused on the rodent-specific, isolation-induced ultrasonic vocalizations of laboratory mice (*Mus musculus*). These studies have found that the PAG^16^ and RAm^18^ both exhibit activity-dependent gene expression during neonatal ultrasonic vocalization, as in adults, as well as identified cell types that appear to play a neonate-specific role in ultrasonic vocalization^24,25^. Unlike laboratory mice, however, parent-directed vocalization in most neonatal mammals, including humans, consists of rhythmic, low-frequency cries produced by the laryngeal vocal folds, rather than ultrasonic vocalizations, which are produced by separate laryngeal mechanisms^26^, and – potentially – separate neural pathways^27^. As a result, the identity of neural pathways that support rhythmic, low-frequency neonatal cries, and the genetic mechanisms by which they have diverged to produce adaptive, species-specific features, remain largely unknown.

Here, we address this gap by examining the neonatal vocal behaviors of deer mice (genus *Peromyscus*), a group of behaviorally diverse, recently diverged North American rodents that, unlike laboratory mice, produce abundant isolation-induced neonatal cries^28^. We find that the duration of cries – a feature associated with infant distress in humans^11,12^ – varies between deer mouse species with distinct social systems, and that, within-species, longer cries elicit faster parental approach. Next, motivated by prior findings in adults, we test the hypothesis that the cerebellar system contributes to interspecies variation in cry duration. Consistent with this hypothesis, we find localized neuroanatomical differences in this brain structure between species with different average cry durations, and show that pharmacological perturbation of the olivocerebellar system alters the duration of cries but not ultrasonic vocalizations. Finally, we use an unbiased forward genetic cross between interfertile deer mouse species to identify loci associated with species divergence in neonatal crying. We find a single quantitative trait locus associated with cry duration on deer mouse chromosome 2, and further identify candidate genes in this locus that are differentially expressed between species, function in early postnatal cerebellar development, and whose loss-of-function phenotype in house mice includes shortened neonatal vocalizations. These findings implicate the cerebellar system in the temporal structuring of neonatal crying and provide a foundation to test molecular and neural hypotheses about the evolution of infant crying in mammals.

## Results

### Variation in temporal features of neonatal cries in recently diverged deer mouse lineages

Across species, parental responses to neonatal vocal signals are shaped by two broad categories of acoustic features^29^: spectral features (e.g., pitch, roughness, and frequency modulation) that reflect how signal energy is distributed in the frequency domain, and temporal features (e.g., duration, rhythm) that describe its distribution in time. Understanding how these feature categories evolve with respect to one another remains an open question in bioacoustics. To address this question in deer mice, we examined previously published neonatal (“pup”) isolation calls recorded from eight deer mouse lineages belonging to four species, *P. maniculatus*, *P. polionotus*, *P. leucopus*, and *P. gossypinus* (Figure 1A). These lineages are recently diverged and encompass variation in habitat (forest vs prairie)^30^, social system (monogamy vs polygamy)^3^, and parental care strategies (mono-parental vs bi-parental)^31^. We have previously shown that isolation calls produced by pups in these species fall into one of two acoustic categories, low-frequency cries (Figure 1B, left) and ultrasonic vocalizations (USVs, Figure 1B, right), with distinct developmental ontogenies, genetic architectures, and functional consequences for eliciting parental behaviors^28^ (cries elicit faster parental approach than USVs). Exploring the acoustic space occupied by these two call types, we noticed they also differ in the variability of their acoustic features, with cries exhibiting qualitatively more variation in acoustic space compared to USVs (Figure 1C). We asked how this variation is partitioned among species by examining 14 spectral and temporal features of pup calls, using species means for each feature to calculate coefficients of variation. We found that temporal features, in particular measures of call duration, were more variable across species than spectral features, a trend that was most pronounced for cries (Figure 1D). Thus, among these deer mouse species, cries exhibit greater interspecies acoustic variation than USVs, and temporal features appear to contribute more to this variation than spectral features.

**Figure 1.**
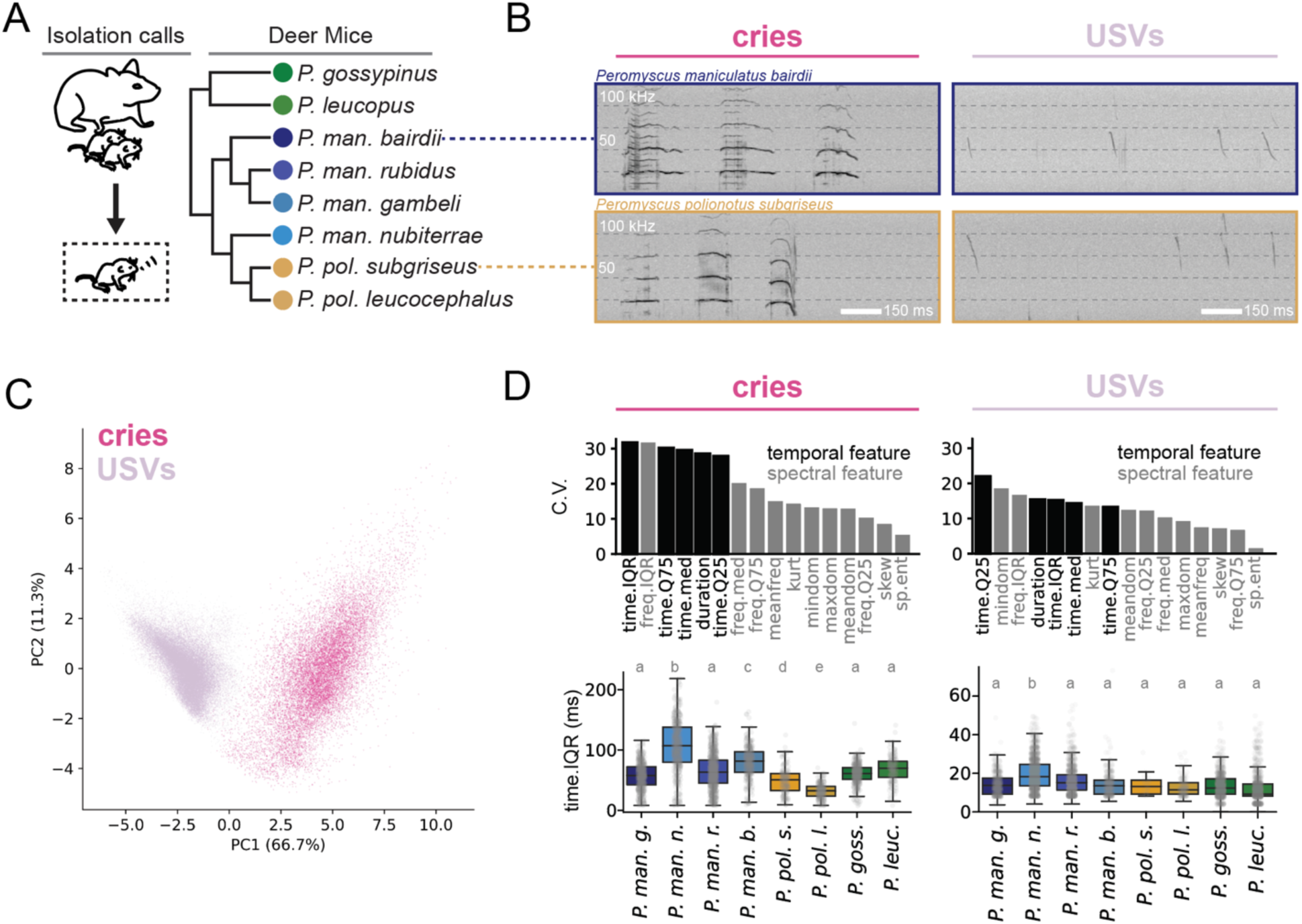
Natural variation in the temporal features of neonatal deer mouse cries. **A.** Left: schematic of protocol for recording isolation calls. Right: Deer mice (genus *Peromyscus*) recorded previously^4^. **B.** Example cries (left) and USVs (right) from two species: *P. maniculatus bairdii* and *P. polionotus subgriseus*. **C.** PCA of acoustic features of all cries and USVs previously recorded from species in panel A, illustrating qualitatively more variability in cries (magenta) than USVs (lavender). **D.** Top left: coefficients of variation of cry acoustic features used for PCA in panel C, calculated from the species mean for each feature. Temporal features of cries are qualitatively more variable than spectral features. Bottom left: A robust measure of call duration (interquartile range of signal energy in time) across species. One-way ANOVA with Tukey post-hoc test (cry: p < 0.001, F = 218.0; USV: p < 0.001, F = 146.7), letters indicate significantly different groups. Right, top and bottom: equivalent analyses from panel D, left, but for USVs.

### The duration of neonatal deer mouse cries affects the speed of parental responses

Motivated by this observation, we examined the potential consequences of cry temporal features on the ability of pups to elicit parental attention. We focused on one temporal feature – cry duration – because in humans^12^ and marmots^11^ longer neonatal cries trigger more rapid parental responses, suggesting that parents interpret cry duration as a signal of need intensity. To test whether cry duration affects parental responses in deer mice, we performed a playback experiment in which *P. polionotus subgriseus* (*P. p. subgriseus*) dams with pups were presented with playback of one of three types of cries (Figure 2A). One of these was unaltered and chosen because its duration was near the *P. p. subgriseus* species mean (120 ms). The other two were artificial manipulations of this cry’s duration while preserving its frequency content, either stretching to a duration approximately two standard deviations above the species mean (“long”, 214 ms) or compressing it by approximately 50% (“short”, 66 ms). All cries were sufficient to elicit behavioral responses from dams (Figure 2B), which included leaving the nest, orienting toward the sound source, and scratching at the cage wall immediately in front of the speaker. However, these responses depended on cry duration. Dams took a qualitatively straighter path toward playback of long cries compared to short cries (Figure 2C, example traces of dam position shown from one trial). They were also more likely to leave their nest in response to long cries than short cries (Figure 2D, paired t-test, t=-2.9, p <0.05), and when they did, they took less time to reach the source of the sound (Figure 2E, paired t-test, t=-3.5, p <0.05). These data suggest that, like humans and marmots, parental *P. p. subgriseus* use temporal information when responding to the sound of isolated neonates, and that longer cries are better able to elicit approach than shorter cries.

**Figure 2.**
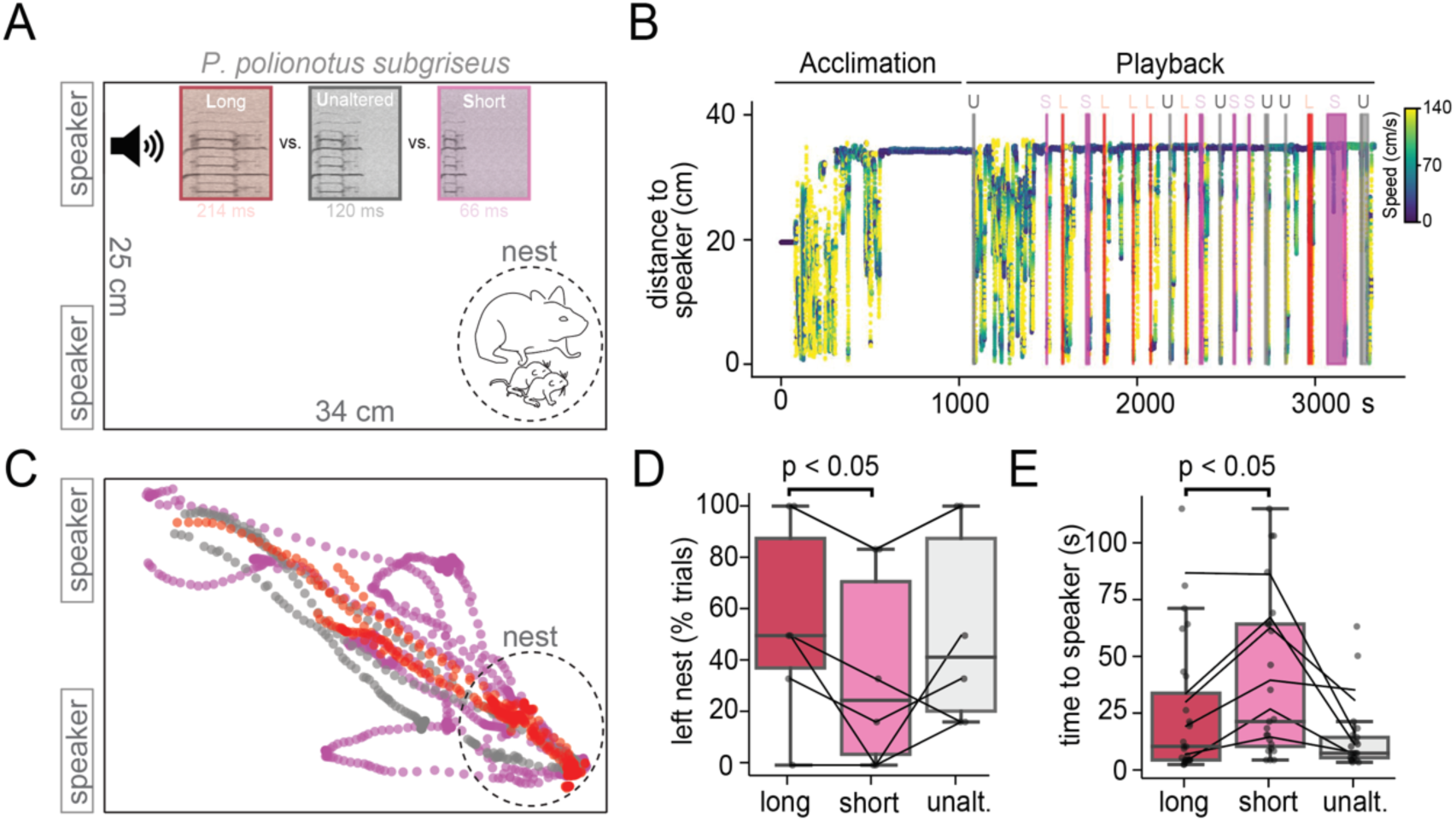
Duration of pup cries affects dam responses in a playback assay. **A.** Schematic of the playback experiment. *P. polionotus subgriseus* dams sitting on litters of postnatal day nine pups were presented with a cry from a postnatal day nine pup of their own species that was either unaltered (120 ms), stretched to 214 ms, or compressed to 66 ms. **B.** Example playback session showing an acclimation phase followed by playback of unaltered (U), short (S), or long (L) cries presented in a randomized order. Y-axis is distance of the dam to the speaker playing the sound. Color represents the dam’s velocity (cm/s). **C.** Example traces of dam position during playback of unaltered (grey), long (red), or short (magenta) cries. **D.** Percent of dams that left their nest in response to playback. More dams leave the nest within 30 seconds following playback of long cries compared to short cries, t = −2.9, p < 0.05, n = 6 dams, paired t-test comparing playback of long and short calls. **E.** Time for the dam to reach within 1 cm of the speaker following long, short, or unaltered cry playback. Dams reach the speaker faster following long cries compared to short cries, t = −3.5, p < 0.05, n = 6 dams, paired t-test comparing playback of long and short calls.

### Species that differ in average cry duration differ in gross olivocerebellar anatomy

The duration of *P. polionotus* and *P. maniculatus* pup cries is not affected by neonatal rearing environment^28^, arguing that, while pups may be capable of tuning cry duration to signal their level of need, species differences in average cry duration are not primarily a result of postnatal learning. We therefore considered whether these differences may arise from species-specific features of neural pathways supporting neonatal crying. Mid-and hindbrain pathways for ultrasonic vocalizations have been described in laboratory strains of the house mouse *Mus musculus*^16^, but, in adults, these pathways are at least partially distinct from those that produce low-frequency calls^27^. Moreover, unlike neonatal deer mice^28^, neonatal *Mus musculus* rarely produce rhythmic, low-frequency cries in isolation (although they are capable of low-frequency vocalization in other contexts).

To guide our search for neural pathways underlying species differences in average cry duration, we sought to identify features that distinguish neonatal deer mouse cries from neonatal deer mouse USVs, focusing on features that are unlikely to arise from differences in morphology (e.g., of the larynx). Temporal structure is one such feature: deer mouse cries are produced in bouts of 2-5 calls, within which vocalizations are separated by silent intervals lasting on the order of 100 milliseconds, and between which vocalizations are separated by more variable intervals lasting on the order of seconds to minutes. This results in distributions of inter-cry intervals that are distinctly bi-modal^28^. In contrast, neonatal deer mouse USVs lack a bout-like temporal rhythm, and exhibit unimodal inter-vocal intervals^28^. To identify neural pathways unique to crying behavior, we therefore chose to examine brain structures that function in the generation of precise motor rhythms.

The olivocerebellar system, consisting of the cerebellum and its major source of excitatory input, the inferior olive, is one example of such a circuit^32–34^ that has recently been implicated in a range of social behaviors^35^, including vocalization in bats^36^, songbirds^37^, and house mice^23^. As interspecies variation in cerebellar anatomy has been linked to behavioral differences in some species^38^, we first examined cerebellar and inferior olive anatomy in postnatal day nine *P. m. bairdii* and *P. p. subgriseus* pups. We measured cerebellar perimeter, a readout of folding in the cerebellar cortex, and cross-sectional area, a readout of cerebellum size, at a set of anteroposterior positions chosen so that they could be matched between species (Figure 3A, B). While *P. m. bairdii* and *P. p. subgriseus* did not differ in the cross-sectional area of the cerebellum at any of these positions (Figure 3C, top), they did differ in perimeter at positions that correspond roughly to lobules V-VII (Figure 3C, bottom)^39^. At these positions, the cerebellum of *P. p. subgriseus* was more folded than that of *P. m. bairdii.* Next, we examined the gross anatomy of the inferior olive (Figure 3D) using a molecular marker (FoxP2) that distinguishes neurons in this structure from the surrounding hindbrain^40^. We found that *P. maniculatus* and *P. polionotus* inferior olives were similar in nuclear counts and cross-sectional area along most of their anteroposterior extents, with the exception of the intermediate inferior olive, where *P. polionotus* both contained more neurons and covered a larger cross-sectional area. The inferior olive and cerebellum are both functionally organized along the anteroposterior axis^39,41^, and cerebellar folding has been proposed to contribute to behavioral diversity across species^38^. Thus, while correlational, the anatomical differences we observe between *P. m. bairdii* and *P. p. subgriseus* could plausibly contribute to temporal differences between neonatal cries in these species.

**Figure 3.**
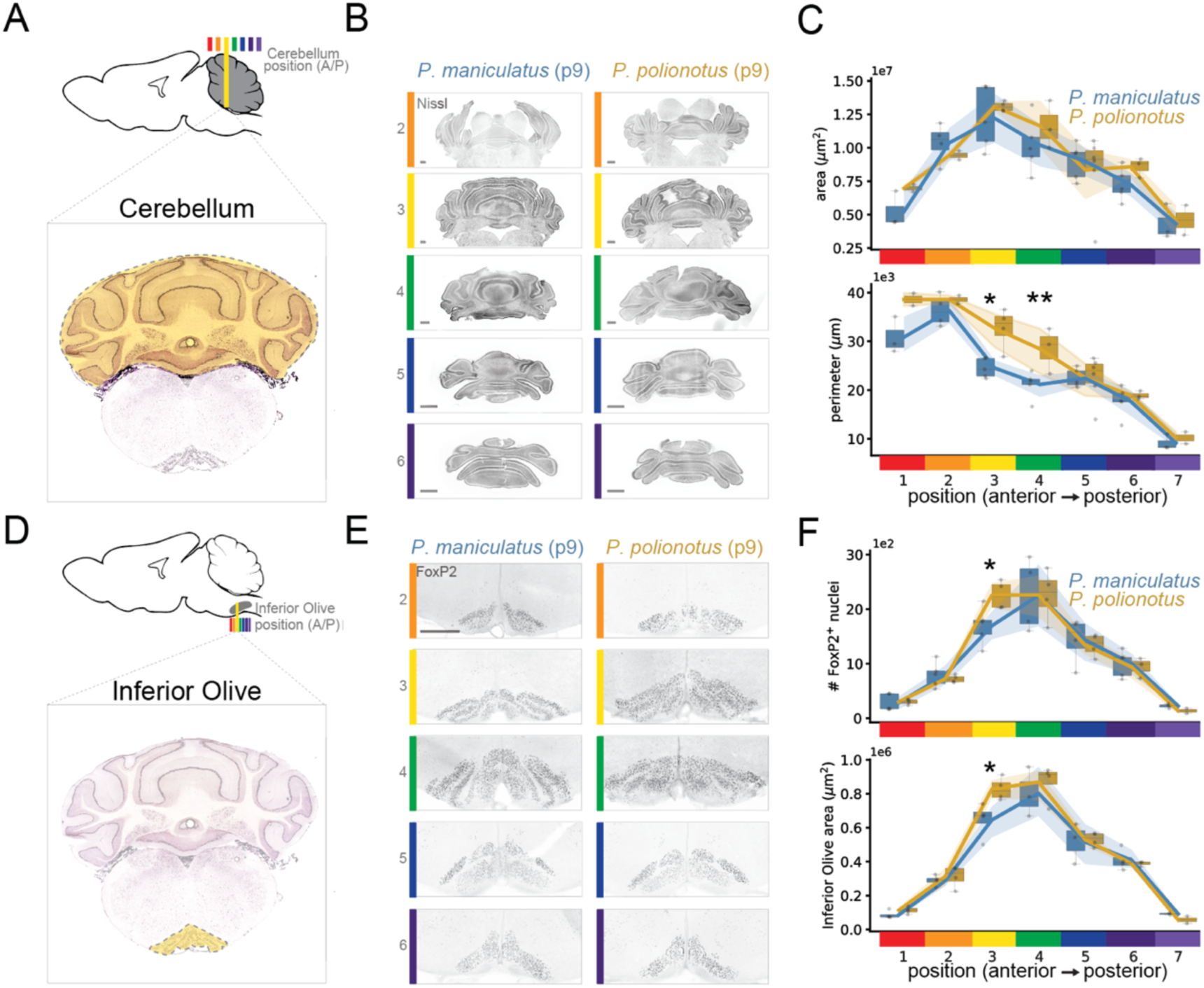
Deer mouse species with different cry durations differ in olivocerebellar anatomy. **A.** Top: schematic mouse cerebellum (grey) with sections taken (colored bars). Bottom: example coronal section of a mouse cerebellum and hindbrain from mouse.brain-map.org/static/atlas. Yellow shading corresponds to the approximate anteroposterior position of the yellow bar above. **B.** Representative Nissl-stained coronal sections from cerebella of postnatal day nine *P. m. bairdii* and *P. p. subgriseus* pups. Colors correspond to bar positions along the anteroposterior axis in panel A, top. Scale bars: 500 µm. **C.** Top: Absolute cross-sectional area of the cerebellum (µm^2^) in postnatal day nine pups at anteroposterior positions illustrated in panels A and B. n.s., not significant. Bottom: Cerebellar perimeter (µm) in the same pups. *P. p. subgriseus* cerebella have larger perimeters in regions 3 (Student’s t-test, t = −2.8, p < 0.05, n = 5 *P. m. b.* and 3 *P. p. s.* pups) and 4 (Student’s t-test, t = −3.9, p < 0.01, n = 5 *P. m. b.* and 3 *P. p. s.* pups). **D.** Top: schematic of the mouse inferior olive (grey) with sections taken (colored bars). Bottom: example coronal section of a mouse cerebellum and hindbrain, yellow shading corresponds to the approximate anteroposterior position of the yellow bar below. **E.** Representative coronal sections from the hindbrain of postnatal day nine pups, stained with a marker of the inferior olive (FoxP2). Colors correspond to bar positions along the anteroposterior axis in panel D, top. Scale bar: 500 µm. All images have the same scale. **F.** Top: Counts of inferior olive (FoxP2^+^) neurons at relative anteroposterior positions illustrated in panels D and E. *P. p. subgriseus* inferior olives contain more FoxP2^+^ neurons at region 3, but not other regions. Student’s t-test, t = −2.9, p < 0.05, n = 4 *P. m. b.* and 4 *P. p. s.* pups. Bottom: Absolute cross-sectional area of the inferior olive (µm^2^) in the same pups. The *P. p. s.* inferior olive is larger at region 3, but not other regions. Student’s t-test, t = −2.7, p < 0.05, n = 5 *P. m. b.* and 4 *P. p. s.* pups.

### Harmaline affects the temporal structure of deer mouse cries

The inferior olive is an intrinsically rhythmic brain region that projects exclusively to the cerebellum, where it is the sole source of the cerebellar climbing fibers^42^. While the function of the inferior olive in behavior has been debated^43^, one model is that its neural rhythms provide the cerebellum with temporal information needed to shape adaptive motor rhythms^44^. Consistent with this hypothesis, manipulating inferior olive activity in anesthetized animals is sufficient to generate motor rhythms in the spinal cord^45^, and in awake animals leads to a loss of motor coordination (“tremor”)^46^. Harmaline^47,48^ is a fast-acting drug that increases the frequency and variability of inferior olive rhythms, generating motor tremor via changes to the conductance of calcium channels in this brain structure^49^, and as such it has long been used as an experimental tool to understand inferior olive function.

To test whether harmaline administration affects the temporal structure of neonatal deer mouse cries, we injected postnatal day nine pups subcutaneously with harmaline or saline, then recorded their isolation-induced vocalizations (Figure 4A). Harmaline but not saline injection increased the duration of pups’ cries by an average of 20% (Figure 4B, saline: n.s., n = 15, t = −0.6; harmaline: p < 0.001, n = 15, t = −6.3). We did not observe an effect of harmaline on the duration of USVs that pups also produce at this age (Figure 4C, saline: n.s., n = 14, t = 0.8; harmaline: n.s., t = 1.1).

**Figure 4.**
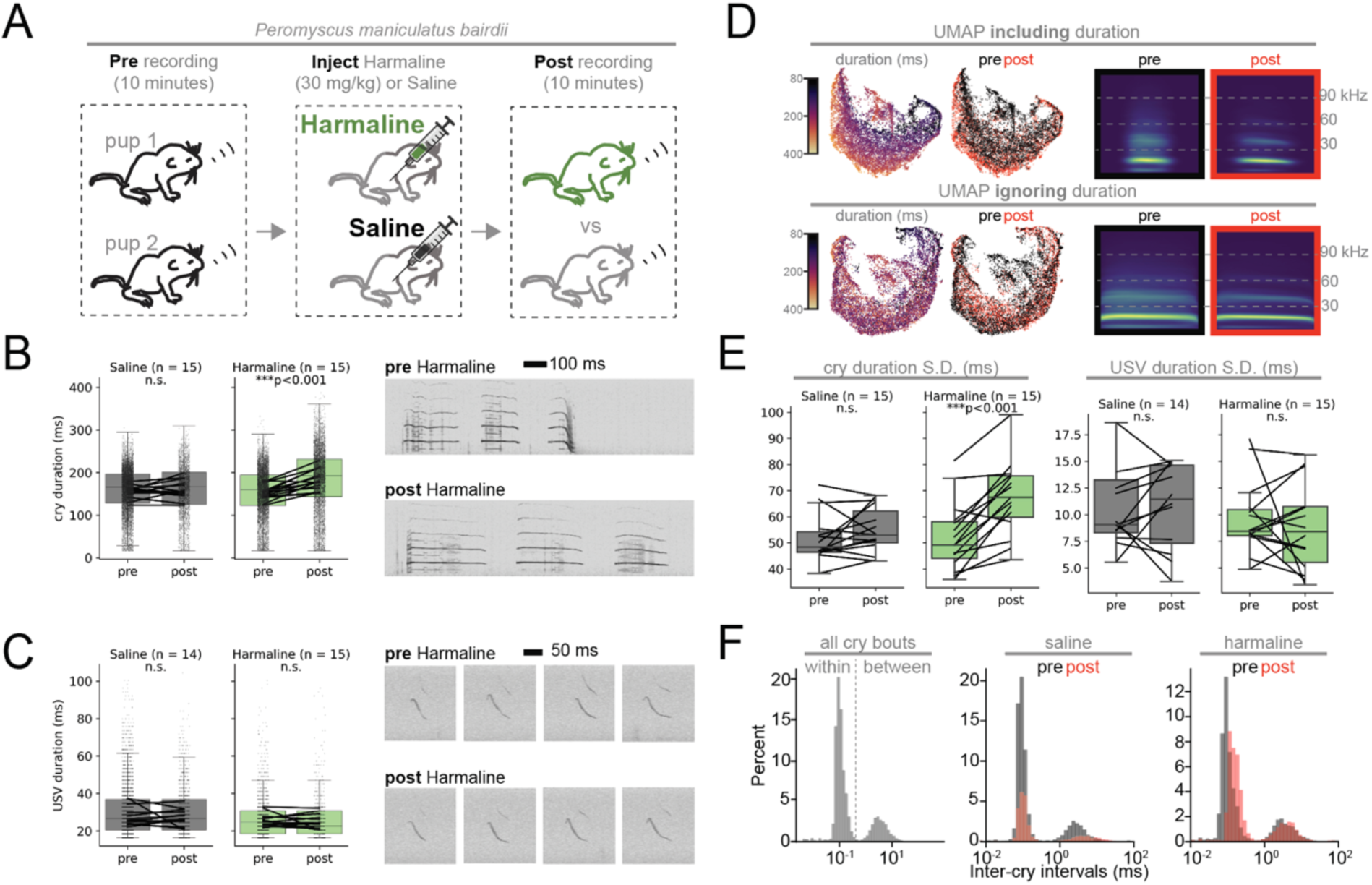
Harmaline alters temporal features of deer mouse cries but not USVs. **A.** Schematic of experimental design. Pups from postnatal day nine *P. m. bairdii* litters were isolated and recorded simultaneously for ten minutes (“Pre”). Pups were injected with 30 mg/kg harmaline or an equivalent volume of saline (“Inject”). Pups were then recorded for another ten minutes in isolation (“Post”). **B.** Left: cry duration in milliseconds, comparing pre- vs post-injection. Saline: grey; harmaline: green. Dots are individual vocalizations. Lines connect averages of each pup. Harmaline significantly increases the duration of cries. Paired t-test, t = −6.25, p < 0.001, n = 15 pups from 6 litters. Right: Example spectrograms of one cry bout from a single pup before (top) or after (bottom) harmaline injection. **C.** Left: USV duration in milliseconds, comparing pre- vs post-injection. Saline: grey; harmaline: green. Harmaline does not affect the duration of USVs. Paired t-test, n.s. Right: Example spectrograms of USVs from a single pup before (top) or after (bottom) harmaline injection. **D.** Top left: UMAP embedding of spectrogram images of all cries, padded to length of longest cry, colored by cry duration (left) or whether the cry was made pre (black) or post (red) harmaline injection (right). Top right: Average padded spectrogram images of all cries recorded pre vs post harmaline injection. Bottom left: UMAP embedding of spectrogram images of all cries, stretched to length of longest cry, colored by cry duration (left) or whether the cry was made pre (black) or post (red) harmaline injection (right). Bottom right: Average stretched spectrogram images of all cries recorded pre- or post-harmaline injection. **E.** Left: Harmaline (right, green) but not saline (left, grey) increases within-pup variance in cry duration (Paired t-test, t = −5.9, p < 0.001, n = 15 pups from 6 litters). Right: Same analysis as left for USV. No effect of harmaline or saline on within-pup variance in USV duration (Paired t-tests, n.s.). **F.** Distributions of inter-cry intervals in all vocalizations (left), pre- or post-saline (middle), or pre- or post-harmaline (right). A threshold of 500 ms (dashed vertical line) was chosen to separate within-from between-bout intervals on the basis of the distribution shown in panel F, left. Harmaline alters the within-bout inter-cry interval (Paired t-test comparing within-pup averages pre- or post-harmaline injection, t = −5.6, p < 0.001), but not the between-bout inter-cry interval (Paired t-test comparing within pup averages pre- or post-harmaline injection, n.s.).

We noticed that harmaline-injected pups lost more body heat than saline-injected pups during the 10 minute recording period (Supplemental Figure 1A, t-test, p < 0.001, t = 5.5, n = 15 pups per treatment), and that they also produced more cries, but not more USVs, in the 10 minute post-injection recording (Supplemental Figure 1B-E). To test whether this difference in heat loss was sufficient to explain differences in vocal duration between saline and harmaline-injected pups, we carried out a second set of paired audio recordings with the same design depicted in Figure 4A, except that rather than injection, pups were placed in a room temperature beaker (“room temperature pups”), or a beaker sitting on top of ice during the second recording (“ice pups”). Pups placed on ice lost heat faster than room temperature pups, as expected (Supplemental Figure 1F, t-test, p < 0.05, t = 3.4, n = 5 room temperature pups and 6 ice pups). However, in contrast to harmaline injection, there was no effect of this cooling on the duration of their cries (Supplemental Figure 1H). These results argue that the effect of harmaline on cry duration cannot be explained solely by harmaline-associated temperature loss.

To further explore the effect of harmaline on cries, we examined the extent to which it altered vocal features other than duration. UMAP (Uniform Manifold Approximation and Projection)^50^ embeddings of cry spectrograms revealed that pre- and post-harmaline injection cries occupy partially non-overlapping regions of acoustic space (Figure 4D, top left). However, artificially stretching all vocalizations to the same length (thus removing from spectrograms any information about call duration), caused the acoustic space occupied by vocalizations recorded pre and post harmaline to become mixed (Figure 4D, bottom left). This suggests that harmaline primarily affects temporal, rather than spectral features, although we note that post-injection cries tend to exhibit less spectral entropy (Figure 4D, compare pre and post on bottom right). Consistent with this qualitative assessment, random forest classifiers trained using a set of 14 spectral and temporal features were able to distinguish pre- vs post-harmaline cries above chance, and cry duration was the feature most important for correct classification. In contrast, spectral features, not duration, were most important for classification of pre- and post-saline cries (Supplemental Figure 1I).

Harmaline affects both the frequency and variability of neural rhythms in the inferior olive^49,51^. We therefore tested whether harmaline injection altered the variability of cry durations produced by individual pups. Harmaline injection significantly increased within pup variance in cry duration, an effect not observed following saline injection (Figure 4E, left. saline: n.s., n = 15 pups, t = −1.3; harmaline: p < 0.001, n = 15 pups, t = −6.0). Harmaline did not change within pup variability in USV duration (Figure 4E, right).

The olivocerebellar system is expected to shape the overall temporal structure of behavior rather than the duration of individual behavioral events^44^. We therefore asked if the effects of harmaline were restricted to cry duration, or if they also extended to the temporal structure of cry bouts. To do this, we examined distributions of inter-cry intervals within and between bouts (Figure 4F, left). These intervals are bimodal, and were largely unaffected by saline injection (Figure 4F, middle, Supplemental Figure 1J-N. See figure legends for details.). In contrast, harmaline injection lengthened silent intervals within bouts, but not between bouts, by an average fold change of 50% (Figure 4F, right. Supplemental Figure 1J-N. See figure legends for details.). Thus, the effect of harmaline on cry duration likely reflects broader effects on the overall temporal structure of crying bouts beyond those on the duration of individual vocalizations.

### A chromosome 2 locus is linked to species differences in cry duration

Taken together, anatomical and pharmacological evidence suggests a model in which the cerebellum contributes to temporal features of neonatal crying in deer mice, consistent with its previously described roles in the temporal structure of adult behaviors. To complement this targeted brain-region-specific approach, we next sought to identify candidate genetic mechanisms by which variation in the temporal features of cries arises between species.

Specifically, we carried out a forward genetic cross between *P. m. bairdii* and *P. p. subgriseus*, which produced 576 second generation (F2) inter-species recombinants. F2s were previously phenotyped by recording isolation-induced vocalizations at postnatal days seven and nine^28^. We sequenced whole genomes of each recombinant individual at low coverage, then used a probabilistic model to infer genotype at 201,303 informative loci per genome^52^. Finally, we used quantitative trait locus (QTL) mapping to test for significant associations between genotype at these loci and the duration of neonatal vocalizations (Figure 5A). Because most phenotyped traits, including duration, were highly correlated between postnatal days seven and nine (Supplemental Figure 2A), we mapped averages of trait values pooled across these ages.

**Figure 5.**
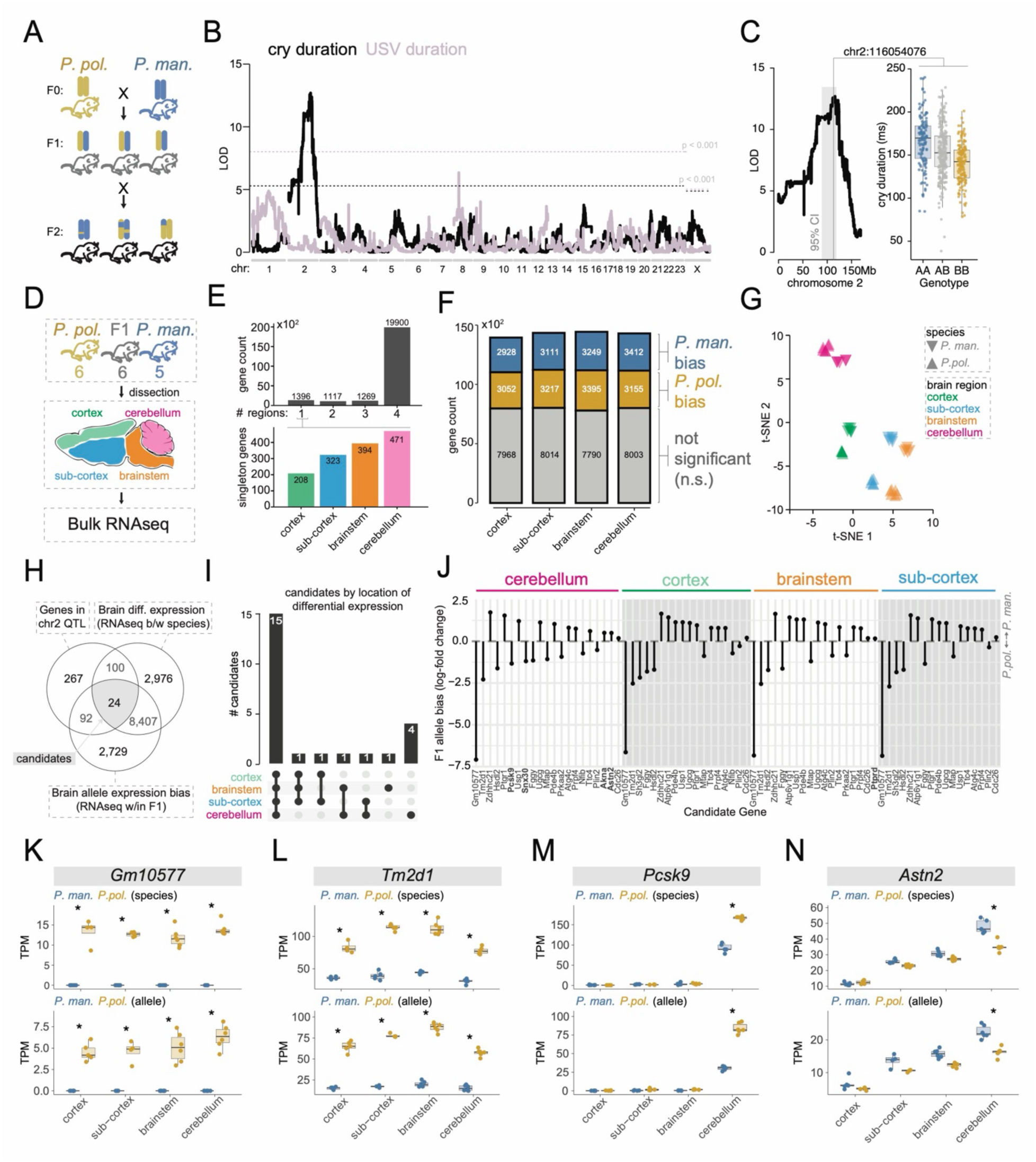
Candidate genes contributing to interspecies variation in neonatal cry duration. **A.** Intercross design. **B.** Quantitative trait loci for cry and USV duration. Genome-wide significance thresholds colored by vocalization type. LOD: log_10_ likelihood ratio, the strength of evidence for a QTL at each marker. **C.** Left: zoom of the chromosome 2 QTL with 95% Bayes credible interval (grey), with the QTL at the location of the vertical grey line. Right: cry duration by genotype at the closest chromosome 2 marker. “A” alleles correspond to *P. man. bairdii,* “B” alleles correspond to *P. pol. subgriseus*. **D.** Neonatal RNA-seq experimental design. **E.** Top: Genes detected in one, two, three, or all four brain regions in both *P. man. bairdii* and *P. pol. subgriseus* samples. Bottom: Of the genes detected in just one brain region, counts by brain region detected. **F.** Total genes detected by brain region (bars) stacked by species bias (blue: *P. man.* bias; tan: *P. pol.* bias). **G.** t-distributed stochastic neighbor embedding of gene expression data labeled by brain region (color) and species (triangle direction). **H.** Intersection strategy to identify 24 genes as candidates to contribute to the chromosome 2 QTL. **I.** Brain regions in which these candidates exhibit cis-differential expression in F1s. **J.** F1 allele bias for all candidates by brain region. Genes with allele bias in just one region are bolded. **K-N.** Transcripts per million by brain region comparing species (top row) or F1 alleles (bottom row) for the two top differentially expressed candidates (*Gm10577* and *Tm2d1*) and two genes differentially expressed in the cerebellum (*Pcsk9* and *Astn2*). Stars indicate brain regions where differential expression was significant both between species and between alleles in F1s.

Consistent with previous work examining trait correlations between cries and USVs in this F2 population^28^, we find that the genetic architecture underlying the duration of neonatal deer mouse cries and USVs is qualitatively distinct (Figure 5B). In addition, we found a single QTL on chromosome 2 that is significantly associated with the duration of cries, but not USVs. Individuals homozygous for the *P. m. bairdii* allele at this locus produced cries on average 18.6% (26.1 ms) longer than those homozygous for the *P. p. subgriseus* allele (Figure 5C, Supplementary Figure 2C), which explains 10.2% of cry duration variance among F2s and accounts for 50.3% of the difference between species in average cry duration (*P. m. bairdii* species average: 171.6 ms; *P. p. subgriseus* species average: 119.7 ms). To assess the specificity of this QTL for cry duration, we first examined correlations between cry duration and three other features: cry spectral centroid, cry count during a ten-minute audio recording, and pup weight (Supplementary Figure 2B). We found that, while these three traits were strongly correlated with each other, they were only weakly (R^2^ = 0.03-0.05) associated with cry duration. In addition, examining significant QTL associated with cry or USV spectral centroid, vocal count, and pup weight, we find significant loci on chromosomes 1 and X, but not 2. Thus, the chromosome 2 QTL is associated with cry duration but not other neonatal vocal traits.

### Fixed coding differences in genes within the chromosome 2 QTL

The 95% Bayes credible interval around the chromosome 2 QTL spans 44.4 Mb (Figure 5C) and contains 483 protein-coding genes. These genes could contribute to the QTL via species differences in coding sequence with functional consequences for vocalization, or via species differences in expression level.

We first determined which genes in the chromosome 2 QTL differ in protein-coding sequence between *P. m. bairdii* and *P. pol. subgriseus.* Using genomic data^53^ from wild-caught individuals from each of these species (15 *P. m. bairdii* and 15 *P. pol. subgriseus*) we found that four of the 483 protein-coding genes in the chromosome 2 QTL contained fixed single-nucleotide polymorphisms (SNPs) that were likely to alter protein function, either because they introduced a non-synonymous change at a conserved site, or because they introduced a premature stop codon. These genes are *Rasef* (Q142`*`), *Mpdz* (R457C), *Dock7* (S26P), and *Cdk5rap2* (G927R). *Rasef* is a Ras family GTPase that regulates the release of digestive enzymes from the pancreas^54^ and has no described roles in brain or behavior. *Mpdz* encodes a PDZ domain protein that functions in the establishment of cell polarity, the regulation of synaptic strength, and whose mouse knockout displays hearing deficits^55^. *Dock7* encodes a guanine exchange factor that functions in axon formation and neuronal polarization^56^. *Cdk5rap2* is a centrosomal protein that functions in microtubule organization. It is critical for spindle orientation during cortical and cerebellar neurogenesis^57,58^, and is associated with microcephaly and developmental delays in humans^59^. There is also evidence that it has been a target of selection influencing brain size evolution in primates^60^, which makes it a particularly promising candidate for future functional studies in deer mice. However, with the possible exception of *Rasef*, all of these variants could plausibly influence behavior via changes to neural circuit development or function, and to our knowledge none have been functionally tested for their effects on vocal production.

### Differentially expressed candidate genes in the chromosome 2 QTL function in cerebellar development

Next, we identified candidates that could plausibly contribute to the chromosome 2 QTL via a difference in expression level between *P. m. bairdii* and *P. p. subgriseus.* Hypothesizing that the postnatal behavioral differences at days seven and nine would likely result from differential gene expression earlier in development, we collected postnatal day 0 (day of birth) brains from *P. m. bairdii, P. p. subgriseus*, and their first generation (F1) hybrids, performing a rough dissection of each brain into cortex, cerebellum, “brainstem” (consisting of hind and midbrain), and “sub-cortex” (consisting of all remaining brain tissue after cortex, cerebellum, and brainstem were removed, Figure 5D). Consistent with a study examining adult brain gene expression in *P. m. bairdii* and *P. p. subgriseus*^61^, the vast majority of mapped genes are expressed in all brain regions examined, with a minority expressed in a single region (Figure 5E, top), of which most were exclusively expressed in the cerebellum (Figure 5E, bottom). Examining differential expression we find – again consistent with adult brain gene expression in these species – that approximately half of neonatal brain expressed genes are differentially expressed between species, with approximately equal numbers of these being biased in expression toward *P. m. bairdii* vs *P. p. subgriseus* (Figure 5F). Finally, we compared samples across species and brain regions using t-stochastic neighbor embedding (t-SNE, Figure 5G). As expected, samples exhibited brain-region-specific expression patterns that differed between species, but were otherwise tightly clustered by brain-region.

We next used this gene expression dataset to narrow down the list of differentially expressed genes to those that could plausibly contribute to the chromosome 2 QTL and thus an interspecies difference in cry duration. Of the 483 genes in the QTL, 100 were significantly differentially expressed between species in at least one brain region. Of these, 24 also exhibited allele-specific expression in F1 hybrids, consistent with the existence of a *cis-*acting mutation within the chromosome 2 QTL that is driving expression differences. The majority (15) of these 24 candidates exhibited both parental and allele-specific differential expression in all four brain regions examined. Four were differentially expressed only in the cerebellum, one only in the brainstem, and the remainder in various combinations of the four regions (Figure 5H, I). Overall, these 24 candidates were significantly enriched for genes differentially expressed in the cerebellum (One-sided Fisher’s Exact Test for cerebellar differential expression among candidates vs all differentially expressed genes: P < 0.01, odds ratio = 4.87. One-sided Fisher’s Exact Test for cerebellar differential expression among candidates vs differentially expressed genes in the chromosome 2 QTL: P < 0.01, odds ratio = 5.56).

To further explore our candidate gene list, we sorted them by magnitude of allelic bias in F1 brains (Figure 5J) and performed a literature review of proposed cellular function and behavioral relevance. This search highlighted four candidates: the top two differentially expressed genes by allelic bias magnitude, *Gm10577* (Figure 5K) and *Tm2d1* (Figure 5L), both of which are differentially expressed in all brain regions, and two genes significantly differentially expressed only in the cerebellum, *Pcsk9* (Figure 5M) and *Astn2* (Figure 5N). Interestingly, it also revealed that one candidate containing a fixed protein-coding difference, *Cdk5rap2*, is strongly enriched in the neonatal cerebellum, although the level of differential expression between species-specific alleles in F1s did not pass genome-wide significance (Supplemental Figure 3).

*Gm10577* is an uncharacterized non-coding gene with no known function that is expressed in neither neonatal (this dataset), nor adult *P. m. bairdii* brains^61^, although it is expressed in both neonatal and adult *P. p. subgriseus*. *Tm2d1* is a beta-amyloid peptide-binding protein implicated in Alzheimer’s Disease^62^ whose *Drosophila* homolog regulates neurogenesis via an interaction with notch signaling. To our knowledge, phenotypes associated with its loss or gain of function have not been described in mammals. *Pcsk9* is a protease with effects on fatty acid metabolism via degradation of low-density lipoprotein receptors^63^. In the house mouse brain, it promotes apoptosis of granule cells in the developing cerebellum^64^, but its functional role in behavior remains debated^65,66^.

*Astn2* is a vertebrate specific transmembrane protein expressed mostly, but not exclusively, in the cerebellum, and is associated in humans with intellectual disability, speech and language delays, and autism spectrum disorders^67^. In the developing cerebellum of *Mus musculus*, it binds to and regulates trafficking of synaptic proteins^67^, as well as the neuron-glial adhesion protein ASTN1, involved in glial guided granule cell migration^68^. *Astn2* knock-out *Mus* exhibit altered Purkinje Cell spine density and corresponding effects on Purkinje Cell synaptic strength^69^. As pups, *Astn2* knock-out animals also produce isolation-induced ultrasonic vocalizations that are both shorter in duration and less numerous than those of wild type pups^69^. While a direct comparison of isolation calls between *Mus* and *Peromyscus* is not possible^28^ (as Mus only produce isolation-induced USVs, not cries), these phenotypes highlight *Astn2* as a candidate for future functional studies in deer mice.

## Discussion

Social environments can create selection pressures that drive behavioral evolution^70^. The neonatal social environment is expected to exert exceptionally strong selection on behaviors that promote infant health and survival, especially in altricial species like mice and humans. Here, we examine neonatal cries in deer mouse species differing in parental care behaviors that may exert such pressures on pup survival. Consistent with this possibility, we observe a correlation between expected parental care levels and the duration of pup cries: pups from promiscuous, mono-parental species with large litters of five to eight pups produce cries that are on average longer than those from monogamous, bi-parental species with smaller litters of two to five pups. We hypothesize that either lower overall levels of parental care, or increased sibling competition, are selection pressures for longer duration cries that are better able to elicit care. However, our phylogenetic comparisons here are limited by a lack of independent contrasts, and we note that there is variation in cry duration between species that do not have reported differences in parental care strategies (e.g., *P. maniculatus bairdii* and *P. maniculatus nubiterrae*). Expanding pup vocal recordings and quantitative measures of parenting to more species will thus be an important next step for quantifying the relationship between neonatal social environments and features of infant crying.

What do the features of infant cries communicate to parents? In the context of adult vocalizations, call timing appears to play a conserved role in signaling affective state, with longer or more rapid calls correlating with heightened arousal across species^71–73^. In the context of neonatal calls, cry duration is associated with parental response in humans^12^ and marmots^11^ (and predator responses in crocodilians detecting distress in the cries of human infants^74^). Here, we find a similar relationship between the duration of individual deer mouse pup cries and the speed of parental responses. This suggests that parental deer mice interpret cry duration as a signal of the magnitude of pup distress. In isolated pup recordings, we nonetheless observe both within-species and within-pup variation in cry duration, even though pups should be experiencing similar levels of distress^28^. Difficult to control features of the nest environment likely contribute to this variation, such as how recently pups fed, their location in the nest relative to others, and litter size, all of which may influence stress-levels. Another possibility is that differences in pup cry duration reflect intrinsic differences in their response to the stress of isolation. Expanding playback experiments to include additional cry-exemplars, species, and environmental (e.g. cold) and physiological (e.g. hunger) triggers will be important for disentangling these possibilities.

Rhythmic, low-frequency cries are a pervasive feature of neonatal social behavior across mammals^29^. Yet the proximate mechanisms supporting neonatal vocalization have been elucidated primarily in mouse genetic models^25^ producing ultrasonic vocalizations. As a result, the identity and function of neural pathways specific to infant crying remain under-explored. Here, we identify the olivocerebellar system as a candidate circuit contributing to temporal features of this behavior. In addition to roles in motor learning^75^, the cerebellum and inferior olive are fundamental to motor timing and temporal pattern generation^33,34,43^ in vertebrates, in some species also contributing to cognitive functions^76^ and social behaviors^35^, including vocalization^37^. On this basis, we test two predictions about the contribution of the olivocerebellar system to neonatal cries: first, that species with different cry durations should differ in olivocerebellar anatomy, features of which (e.g., foliation) have been linked to variation in species-specific motor patterns^38,77^, and second, that systemically perturbing olivocerebellar activity should lead to corresponding alterations in the temporal structure of pup cries. Examining olivocerebellar anatomy, we observe a correlation between temporal features of cries and anatomical features of the cerebellum (foliation) and inferior olive (size, number of neurons) that are localized to specific regions along their anteroposterior axes. Furthermore, harmaline, a drug that alters the rate and variance of neural rhythms in the olivocerebellar system, alters the mean duration and variability of neonatal cries, but not ultrasonic vocalizations.

These findings are consistent with a role for the olivocerebellar system in the generation of neonatal cries, but our methodologies come with limitations. First, we examined gross cerebellar and inferior olive anatomy in two species; examining the molecular features and micro-architecture of these brain structures across all the species for which we have neonatal vocal data will be important for testing the relationship between neuroanatomy and vocal behavior we propose here. In addition, we have used a pharmacological approach (subcutaneous injection) that is likely to produce off-target effects on behavior. We believe the specificity of the harmaline effect to cries (but not USVs) is a compelling indication that these off-target effects are not dominating our behavioral data, but a more targeted approach (e.g., micro-injection, optogenetics) is needed to causally link this region to temporal features of crying. Finally, we note that we have focused here on neural mechanisms because the temporal structure of crying is likely under neural (rather than morphological) control. However, laryngeal morphology may be more important than neural mechanisms for explaining other features of infant cries. Understanding how variation in the larynx relates to species variation in these features is a promising direction for future research.

Even if targeted experimental manipulations reveal a functional connection between the cerebellum and vocal production circuits in neonatal deer mice, as they recently have in adult laboratory mice^23^, it leaves open the question whether naturally occurring species differences in the cerebellum or other brain regions cause meaningful differences in neonatal vocal behavior. Forward genetics provides one way to answer this question. It also provides an independent entry point to discover contributions to vocal behaviors without making assumptions about underlying mechanism. While interspecies genetic mapping is not an option in most experimental systems due to reproductive barriers, deer mice are an exception in which some behaviorally divergent species remain interfertile in lab settings. We take advantage of this feature here to perform an intercross between two species, *P. m. bairdii* and *P. p. subgriseus*, differing in the duration of their neonatal isolation cries, and find a 44 Mb locus on chromosome 2 associated with cry duration.

Of the 483 genes in this locus, four contained fixed non-synonymous substitutions with moderate to high predicted impact on protein function. Three of these are expressed in the neonatal deer mouse brain (*Mpdz*, *Dock7*, *Cdk5rap2*) and could plausibly affect neonatal vocal behaviors. *Cdk5rap2* is particularly interesting. It contains the most non-synonymous SNPs of any gene in the chromosome 2 QTL (5, although only one is in a highly conserved site), humans carrying *Cdk5rap2* mutations have altered cortical and cerebellar volumes^59^, and in primates it appears to be under selection for its effects on brain size^60^. Moreover, in our neonatal brain RNA-seq dataset, *Cdk5rap2* is enriched in the cerebellum and is differentially expressed between species, although the level of differential expression did not pass genome-wide significance in F1s. In addition to *Cdk5rap2*, 24 genes (none of which contain fixed coding variants) are differentially expressed in the brains of neonatal *P. m. bairdii* and *P. p. subgriseus* and their F1s in a manner that could plausibly contribute to the chromosome 2 QTL. Four of these genes are only differentially expressed in the cerebellum. Of these, *Astn2* is particularly interesting. *Astn2* is a gene implicated in autism spectrum disorders that include language delays, and functions in cerebellar development by facilitating glial-mediated migration of granule cells, as well as Purkinje Cell synaptic physiology^67,69^. Moreover, *Astn2* knock-out lab mice (*Mus musculus*) produce shorter and fewer isolation-induced neonatal USVs^69^. Neonatal *Mus* do not produce isolation-induced cries, so a comparison to the low-frequency cries of neonatal deer mice should be treated with caution. Nonetheless, the *Astn2* mutant phenotype resembles the naturally occurring phenotypic difference we find between *P. man. bairdii* and *P. pol. subgriseus*: *P. man. bairdii* expresses *Astn2* at a higher level than *P. pol. subgriseus* in the cerebellum, and also produces longer and more numerous^28^ cries. *Astn2* and *Cdk5rap2* are therefore both compelling candidates for future studies of neonatal vocal behavior evolution.

Together, our findings suggest a hypothesis for the evolution of neonatal vocal repertoires in which changes in the expression level (e.g. of *Astn2*) and/or function (e.g., of *Cdk5rap2*) of genes regulating cerebellar development lead to differences in cerebellar anatomy and/or physiology, which in turn lead to differences in temporal features of neonatal cries that alter the probability of parental approach. Future studies should functionally manipulate *Astn2, Cdk5rap2,* and the other candidate genes identified here to test these causal hypotheses in deer mice.

## Experimental model and subject details

Data in Figure 1 were generated using eight *Peromyscus* taxa, representing four species (*P. maniculatus*, *P. polionotus*, *P. leucopus*, *P. gossypinus*), as described previously^28^. All animals were housed in barrier specific, pathogen-free conditions with 16 h light: 8 h dark at 22° C in individually ventilated cages (18.6 cm x 29.8 cm x 12.8 cm height; Allentown, New Jersey) with quarter-inch Bed-o-cob bedding (The Andersons, Maumee, Ohio). Breeding animals and their litters were fed irradiated PicoLab Mouse Diet 20 5058 (LabDiet, St. Louis, Missouri) *ad libitum* and had free access to water. We weaned animals at 23 days of age into same strain and sex cages. After weaning, we fed animals irradiated LabDiet Prolab Isopro RMH 3000 5P75 (LabDiet) *ad libitum* with free access to water and provided them with nesting material (Nestlet, Ancare, Bellmore New York) and a polycarbonate translucent red hut. All experiments were approved by the Harvard University Faculty of Arts and Sciences Institutional Animal Care and Use Committee.

## Methods details

### Audio recording

Pups were recorded in sound-attenuating chambers with CM16/CMPA ultrasonic microphones (Avisoft, Berlin, Germany) at a 250 kHz sampling rate and 16-bit encoding, as described previously^28^.

### Vocal segmentation and feature extraction

Raw audio recordings were segmented into vocalizations using signal amplitude with the get_onsets_offsets function of the python package AVA^78^ as described previously^28^. Acoustic features of segmented vocalizations were calculated using the R package warbleR^79^.

### UMAP embedding

Spectrogram images of vocalizations were embedded using the fit.transform method of the python package umap-learn and default settings, as described previously^28^. To compare effects of harmaline on duration to effects on other acoustic features, spectrograms were either padded to the length of the longest vocalization with zeros (preserving information about cry duration in the image) or stretched to the length of the longest vocalization (removing information about duration).

### Random forest models

We used the Python package scikit-learn (version 1.0.2) to train random forest models. Models were trained to classify cries as either “pre” or “post” injection. We trained two models (one for harmaline injection, one for saline), each with 100 trees using Gini impurity as the optimization criterion. Features for training were calculated with the R package warbleR^79^ and are as follows (see warbleR documentation for details).

duration: time from start to end of cry
time.ent: entropy of the signal in time
time.Q25: first quartile of signal energy in time
time.Q75: third quartile of signal energy in time
time.IQR: time range between time.Q25 and time.Q75
time.median: time at which signal is divided into two intervals of equal energy
freq.Q25: first quartile of frequency (kHz)
freq.Q75: third quartile of frequency (kHz)
freq.med: median frequency (kHz)
sd: standard deviation of frequency (kHz)
meanfreq: average frequency (kHz)
sfm: spectral flatness
freq.IQR: frequency range between ‘freq.Q25’ and ‘freq.Q75’ (in kHz)
sp.ent: entropy of the signal in frequency domain

### Acoustic playback: data collection

Breeding pairs of *P. polionotus subgriseus* were checked daily for pups. When pups from a given breeding pair were 8 days old, we moved the mother, her nest, and her litter to the cage in which we performed the playback experiments (25 cm x 34 cm x 19 cm height; Allentown, Allentown, New Jersey). We modified the cage to have two grids of holes, on either end of one wall, for audio playback. After 24 hours, we moved the cage containing the mother and her litter to a separate room and left it on a table-top for 10 minutes. Once the mother was in her nest for an additional 1 minute, we played pre-recorded pup cries from one of the two ultrasonic speakers (Avisoft) - on a loop with cries separated by 1s of silence - until the mother touched the cage wall in front of the speaker, or for 2 minutes, whichever came first. We then stopped playing the audio recording until the mother returned to her nest and remained there for 1 minute, at which point we recommenced playback. This regime continued until each playback vocalization was presented six times in a random order determined using the *sample()* function of the python package *random*. We used an ultrasonic microphone placed above the nest to confirm that both vocalization types were detectable at the location of the mother during playback. All playback experiments were conducted in red light within one hour of the onset of the animals’ dark cycle.

### Acoustic playback: analysis

We used the software package bonsai to track the mother’s position during playback and align position measurements to the active speaker^80^. All audio was recorded using the hardware and recording specifications described above to record pup vocalizations. To produce audio for the playback experiment, we first chose a species-typical *P. polionotus subgriseus* cry whose duration (120 ms) approximately matched the species mean, as determined previously^28^. Using Adobe Audition (Adobe Inc.), we then stretched this cry to approximately two standard deviations about the species mean (214 ms) or compressed its duration to about 50% of the species mean (66 ms). Shorter vocalizations were not tested because further shrinking introduced acoustic artefacts. All playback vocalizations were matched in amplitude in Adobe Audition before being exported as 16-bit wav files.

### Tissue Collection for Neuroanatomy

We deeply anaesthetized postnatal day nine *P. maniculatus bairdii* and *P. polionotus subgriseus* pups with a mixture of ketamine and xylazine, then perfused them with 4% paraformaldehyde. Following perfusion, brains were harvested, cryo-protected in a 30% weight by volume solution of sucrose in phosphate-buffered saline (PBS) and stored in embedding medium (Fischer Scientific Tissue Plus O.C.T Compound) at −80 °C until sectioning. 60 µm sections of hindbrain and cerebellum were collected and stored in cold PBS +0.25% triton. Five *P. maniculatus bairdii* and four *P. polionotus subgriseus* pups were sectioned.

### Immunohistochemistry

Immediately following sectioning, free floating brain slices were washed 3 times in PBS +0.25% triton for 20 minutes each at room temperature, then blocked in 3% normal goat serum in PBS +0.25% triton for one hour, also at room temperature. They were then stained with rabbit anti-FoxP2 antibody (Abcam, AB16046) diluted 1:1000 in blocking solution (PBS +0.25% triton + 3% normal goat serum) for 48 hours at 4°C on a shaker. Next, they were washed 3 times in PBS +0.25% triton for 20 minutes each at room temperature, before being stained with donkey anti-rabbit antibody (Thermo Fisher catalogue **#**A-21209) diluted 1:1000 and Fluorescent Nissl stain (ThermoFisher Scientific NeuroTrace, N21482) diluted 1:200 for 3 hours at room temperature. Sections were then mounted with DAPI and imaged on a Zeiss Axio Scan.Z7 slide scanning microscope.

### Comparative anatomy

Prior to analysis, all images were sorted according to anterior/posterior position and aligned between species using the beginning and end of each brain structure as reference points. For each anteroposterior position, damaged sections were excluded. Nissl-stained coronal sections of the cerebellum were traced using the ImageJ free hand drawing tool. Area and perimeter of these outlines were calculated using the ImageJ “measure” function. Anatomical features of the inferior olive were measured using FoxP2 stained coronal hindbrain sections from the same pups that generated sections for anatomical data on the cerebellum. Cross sectional area was calculated in ImageJ, as above, by free hand tracing the perimeter of the inferior olive, as revealed by FoxP2 positive nuclei. FoxP2 positive nuclei were counted using an automated image segmentation algorithm (micro-sam^81^) and validated by a subset of hand-counted samples.

### Pharmacology

Harmaline Hydrochloride Dihydrate was obtained from Indofine Chemical Company (Hillsborough, NJ) and dissolved in sterile physiological saline (0.9% NaCl) to a concentration of 1 mg/mL. Dilutions were made fresh prior to each experiment, which were conducted on litters of postnatal day nine *P. maniculatus bairdii* pups. Entire litters (N = 6) were isolated and recorded simultaneously for 10 minutes, as described previously^28^. During pre-injection recording, pups were randomly selected to receive either a 30 mg/kg subcutaneous injection of harmaline or a subcutaneous injection of an equivalent volume of saline. Following the pre-injection recording, pups were injected with saline or harmaline, then left for 1 minute on a warming pad set to 36°C before being recorded again for 10 minutes. Acoustic data were subsequently segmented into cries and USVs, as described previously^28^, and features of these vocalizations were compared within each pup pre- and post-harmaline or saline injection using averages over vocalizations in each ten-minute recording.

### Genetic Cross Design

We carried out an intercross between *P. maniculatus bairdii* and *P. polionotus subgriseus* as described previously^28^. Briefly, two *P. m. bairdii* females were crossed with two *P. p. subgriseus* males to generate 54 first generation (F1) hybrids, who were then paired to generate 576 second generation (F2) hybrids. The isolation-induced vocalizations of each F2 animal were recorded once on postnatal day seven and once on postnatal day nine. Vocalizations were recorded as described previously^28^. After the postnatal day nine recording, animals were sacrificed, and liver tissue was harvested for sequencing.

### Whole genome sequencing

We performed low-coverage whole genome sequencing of 576 F2 animals for which isolation-induced vocalizations were recorded. For each F2, 50 mg of liver tissue was lysed in a solution containing Tissue Lysis Buffer A, Proteinase K, and RNAse A on a shaker at 56 °C overnight. Genomic DNA was subsequently purified using a Maxwell RSC Tissue DNA Kit (Promega). Sequencing libraries were prepared using the Illumina DNA library preparation workflow with an expected insert size of 200-300 bp. Library quality and fragment size distributions were verified using a 2200 TapeStation (Agilent Technologies). Libraries were sequenced to an average depth of 0.19x (average of 1.6×10^6^ reads per sample) on an Illumina NovaSeq 6000 across two lanes (288 samples per lane) of an SP flow cell, with paired end, 2 x 151 bp reads and dual 10bp indexes. Raw base call files were converted to FASTQ and demultiplexed (assigned to samples) based on exact matches to the 10 bp index sequences using Illumina’s GenerateFASTQ workflow. Reads were mapped to the *P. maniculatus bairdii* genome (GenBank assembly accession GCA_003704035.3) using the bwa-mem algorithm from the software package BWA (version 0.7.17)^82^. Genotype (ancestry) probabilities were inferred using the fitHMM method of the Multiplex Shotgun Sequencing (MSG) package (version v0.4.10) and filtered to informative sites with the following parameters in the MSG pullthin script:

deltapar1 = 0.1
deltapar2 = 0.1
recrate = 25
rfac = 1
priors = {0.25, 0.5, 0.25}
theta = 1

This resulted in 201303 markers used for mapping. The genotype probabilities at each site were passed to a cross object in the R package qtl (v1.60) using the read.cross.ms g function of MSG.

### Quantitative Trait Locus Mapping

We performed quantitative trait locus mapping using Haley-Knott regression implemented by the scanone function of R/qtl (v1.60). Because we found that cry duration was highly correlated between postnatal day seven and postnatal day nine recordings, we mapped the average duration of cries (or USVs) made by each pup in each recording. Genome-wide significance thresholds were calculated on the basis of 1000 genotype-phenotype permutations, implemented by scanone in R/qtl with the n.perm parameter set to 1000. 95% Bayesian credible intervals were calculated for significant peaks using the bayesint function in R/qtl. Percent variation explained by the chromosome 2 QTL was calculated using the formula 1 - 10^−2lod/n^, implemented by fitqtl in R/qtl.

### Brain RNA-seq

We collected brain tissue from postnatal day 0 mouse pups from *P. m. bairdii, P. p. subgriseus*, and their first-generation hybrids. 9 breeding pairs (3 *P. m. bairdii*, 3 *P. p. subgriseus*, 3 containing *P. m. bairdii* females and *P. p. subgriseus* males) were checked daily to ensure all harvested brain tissue came from pups born within the last 24 hours. Pups were euthanized by five minutes of CO2 exposure. Brains were then immediately removed in ice-cold phosphate-buffered saline and dissected into four anatomical regions (cerebellum, cortex, mid/hind brain, and sub-cortex) using a scalpel, then immediately flash frozen in liquid nitrogen.

Total RNA was isolated using an RNeasy 96 QIAcube HT Kit (Qiagen). mRNA was subsequently isolated using polyT oligo-attached magnetic beads, then used to generate complementary DNA (cDNA) libraries. Libraries were sequenced on an Illumina NovaSeq X Plus (paired end, 2x 150 bp). Library prep and sequencing were carried out by Novogene, Inc. (California, USA). Raw data was filtered by removing reads containing adapters, removing reads where >10% of bases could not be determined, and >50% of bases contained a quality score (Qscore <=5). Remaining reads were further filtered using trimmomatic (version 0.39) by removing 5’ and 3’ ends until a Qscore >= 3, and by dropping reads shorter than 36 base pairs. Cleaned, trimmed reads were mapped to their corresponding reference genomes (*P. m.* bairdii, GenBank assembly accession GCA_003704035.3, and *P. pol.* subgriseus, GenBank Accession GCA_003704135.2), and custom annotations generated with *Comparative Annotation Toolkit*^83^. F1 samples were mapped to a diploid reference genome generated by concatenating the *P. m. bairdii* and *P. pol. subgriseus* assemblies and annotations. Read mapping and expression estimates were carried out using the rsem-calculate-expression method of the package RSEM (v1.3.1). Mapped genes were filtered to retain only those present in both species’ annotations (n = 33,836).

### PCA and t-SNE

The PCA in Figure 1C was performed using scikit-learn (sklearn.decomposition.PCA) on centered, scaled acoustic features of vocal segments published previously^28^. Raw data to reproduce these features can be found here: https://doi.org/10.5061/dryad.g79cnp5ts. The t-SNE in Figure 5G was performed on gene counts transformed using variance stabilizing transformation implemented by the vst function of the R package DESeq2. T-distributed stochastic neighbor embedding was performed using scikit-learn (sklearn.manifold.TSNE) with perplexity set to 10.

### Detecting fixed protein-coding differences

To identify fixed coding differences in the chromosome 2 QTL, we examined whole genome sequence data from 15 wild-caught *P. maniculatus bairdii* and 15 *P. polionotus subgriseus* individuals^53^. To be considered a fixed difference, variants had to be called in all individuals of each species, be homozygous in all called individuals, and have different alleles in each species. To identify variants affecting protein sequences, we used a SnpEff annotated vcf (variant call file) to select for variants with moderate or high predicted impact. We then predicted which variants are likely to impact protein function, using MAFFT (v7.310) to generate multiple sequence alignments between *P. m. bairdii* protein sequences and homologs identified in the blast nr database, retaining variants in regions with evidence of evolutionary constraint, defined as conservation of the reference amino acid in >80% of homologs.

### Differential expression analysis

For each dissection brain region, we tested for differential expression between species using DESeq2 (v1.42.1). Wald tests were applied per gene using count estimates produced by RSEM, with size factors and dispersions estimated using default settings by DESeq2. The Benjamini-Hochberg method was used to correct for multiple comparisons, and genes with an adjusted p-value < 0.05 were considered significantly differentially expressed. Cis-differential expression was assessed in F1s using DESeq2 by comparing allele-specific count estimates generated by mapping reads to the concatenated *P. m. bairdii* and *P. pol. subgriseus* assembly. The Benjamini-Hochberg method was used to correct for multiple comparisons in DESeq2. In addition, we compared transcripts per million (TPM) across brain regions for each gene using ANOVA with a Tukey posthoc test, comparing species or (in F1s) alleles, and accounting for multiple comparisons across brain regions using Bonferroni correction. Of the genes with an adjusted DESeq2 differential expression p-value < 0.05, those that also had a TPM difference p < 0.05 were considered significantly differentially expressed candidates. The final candidate list was generated by the intersection of three lists: the genes differentially expressed between *P. m. bairdii* and *P. pol. subgriseus*, the genes differentially expressed in *cis* (i.e. significant differential expression between allele origin in F1s) in the same direction as in trans (i.e. between species), and genes contained within the 95% Bayes credible interval underneath the chromosome 2 QTL peak.

## Acknowledgments

We would like to thank Jenny Chen and Nacho Sanguinetti-Scheck for providing feedback on this manuscript. We also thank Michael Long, David Schneider, Gregg Castellucci, and Tim Sainburg for suggestions and advice throughout the course of this study.

## Author contributions

N.J. and H.E.H. conceived and planned the project. N.J. and M.L.W. collected and analyzed the data. N.J., M.L.W. and H.E.H. wrote the manuscript.

---

**Supplemental Figure 1.**
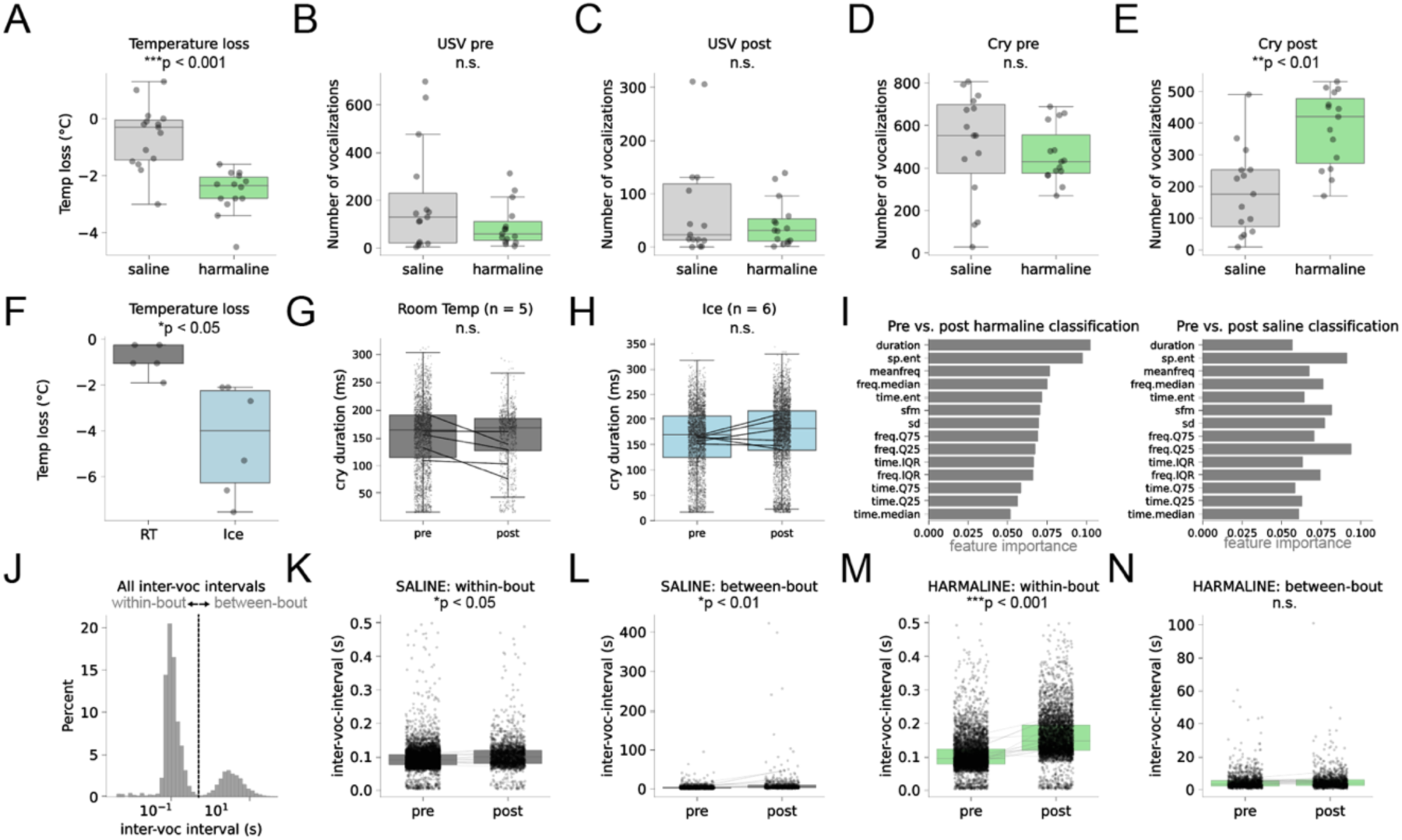
Harmaline alters neonatal cries independent of temperature loss. **A.** Pups lost more heat in 10 minutes following harmaline vs saline injection (p < 0.001, t = 5.3, n = 15 pups per condition). **B-E.** Cry and USV counts during 10-minute recordings pre-injection (pre) or post-injection (post) of saline and harmaline. Harmaline-injected pups cry more than saline-injected pups (p < 0.01, t = −4.2, n = 15 pups per condition). Harmaline does not affect the number of USVs pups produce (C, n.s., n = 15 pups per condition). Harmaline- and saline-injected pups vocalize at similar rates prior to injection (B, D, n.s., n = 15 pups per condition). **F.** Temperature loss of pups recorded at room temperature (RT) or on ice (p < 0.05, t = 3.5, n = 5 RT pups and 6 ice pups). **G**. Recording at room temperature does not change cry duration (paired t-test comparing average duration of pup cries, n.s.). **H.** Recording on ice does not change cry duration (paired t-test comparing average duration of pup cries, n.s.). **I.** Feature importances of random forest classifiers trained to label cries as pre- or post-harmaline (left) or pre- or post-saline (right) injection. **J**. Distribution of all inter-vocalization intervals recorded pre-injection of saline or harmaline (reproduced from Figure 4F, left). Vertical line at 0.5 seconds indicates threshold for distinguishing within-bout intervals from between-bout intervals in panels K-N. **K-N**. Effect of saline and harmaline injection on within- vs between-bout inter-vocalization intervals (paired t-test comparing within pup averages pre vs post-injection, n = 15 pups per plot). Saline mildly affects within-bout intervals (K, p < 0.05, t = −2.4) and between-bout intervals (L, paired t-test comparing within pup averages, p < 0.001, t = −3.6). Harmaline significantly affects within-bout intervals (M, p < 0.001, t = −5.6) but not between-bout intervals (N, n.s.).

**Supplemental Figure 2.**
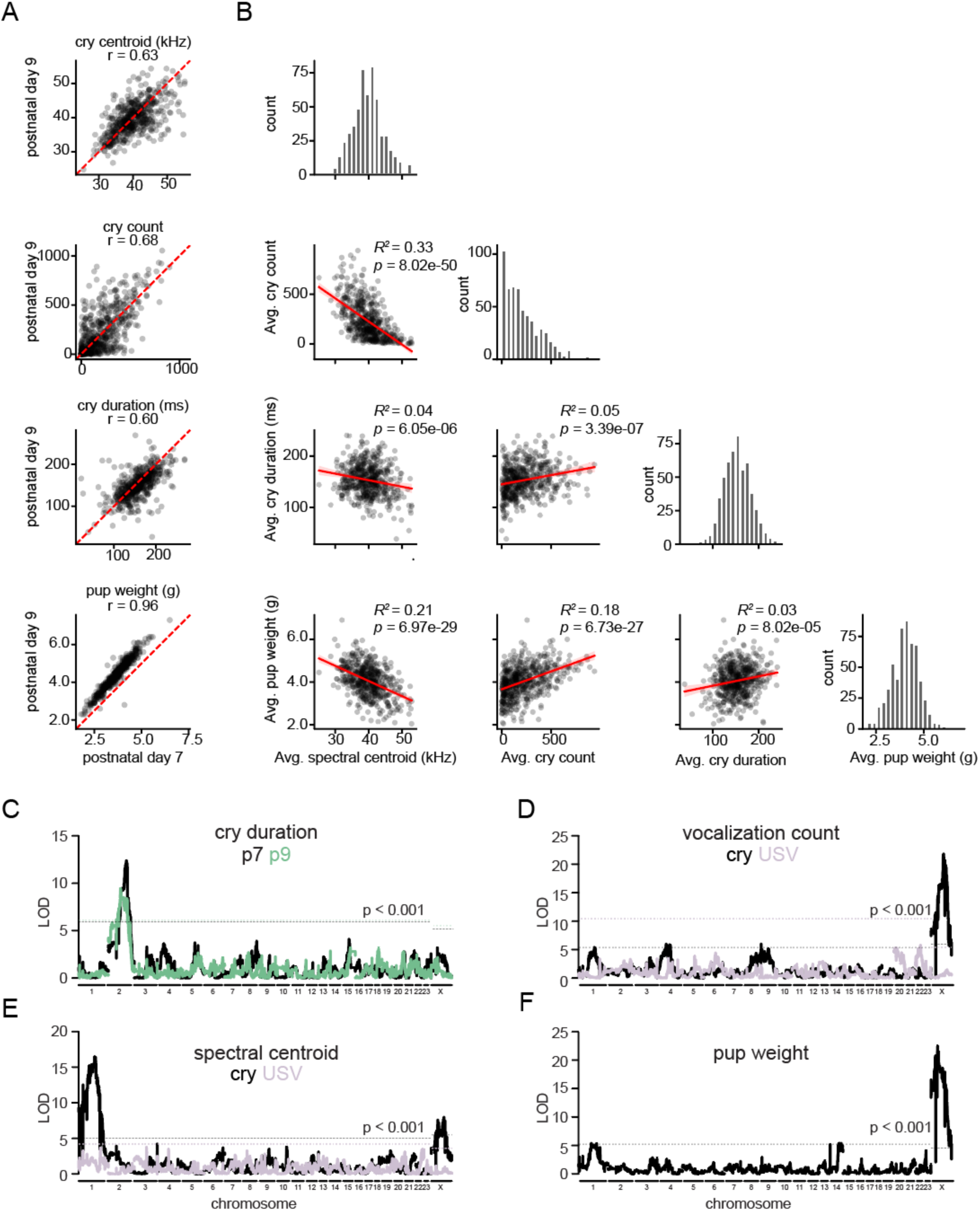
A chromosome 2 QTL is specific to duration but not other features of neonatal cries. **A.** Cry features (spectral centroid, count during 10-minute recording, duration) and weight averaged by F2 pup at postnatal day seven and postnatal day nine. Red dashed line indicates y = x. Pearson correlation coefficients (r) at the top of each plot. **B**. Pairplot quantifying correlations among traits plotted in panel A, using averages of pooled postnatal day seven and postnatal day nine data per pup. **C.** Quantitative trait loci for cry duration, mapped separately at postnatal day seven (black) and nine (green). Genome-wide significance thresholds colored according to age. **D-F**. Quantitative trait loci for spectral centroid of cries or USVs (E), count of cries or USVs in a 10-minute recording (D), and pup weight (F). LOD: log_10_ likelihood ratio.

**Supplemental Figure 3.**
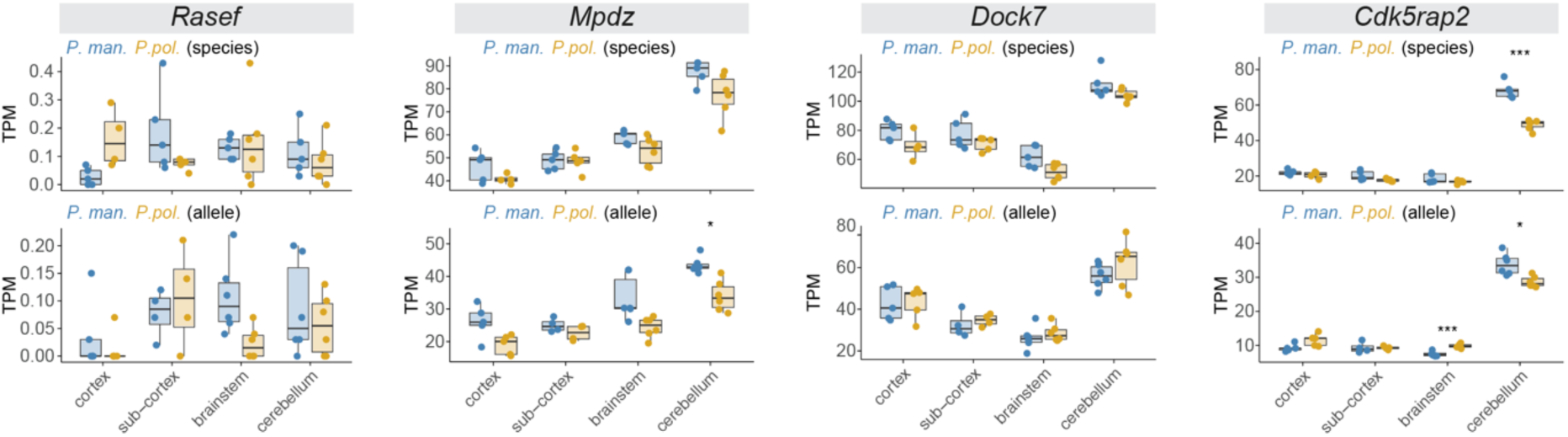
Neonatal brain expression of genes in the chromosome 2 QTL with fixed coding differences between *P. m. bairdii* and *P. p. subgriseus*. TPM: Transcripts per million. *p < 0.05, ***p < 0.001, ANOVA with Tukey posthoc test comparing species (top) or allele (bottom) within brain region. Within each gene, p-values are Bonferroni corrected to account for multiple comparisons across brain regions. No genes pass genome-wide significance for differential expression between species and between alleles.

## Notes

### Competing Interest Statement

The authors have declared no competing interest.

